# LncATLAS database for subcellular localisation of long noncoding RNAs

**DOI:** 10.1101/116335

**Authors:** David Mas-Ponte, Joana Carlevaro-Fita, Emilio Palumbo, Toni Hermoso, Roderic Guigo, Rory Johnson

**Author notes:** Equal contribution (David Mas-Ponte, Joana Carlevaro-Fita).

## Abstract

**Background:** The subcellular localisation of long noncoding RNAs (lncRNAs) holds valuable clues to their molecular function. However, measuring localisation of newly-discovered lncRNAs involves time-consuming and costly experimental methods.

**Results:** We have created “LncATLAS”, a comprehensive resource of lncRNA localisation in human cells based on RNA-sequencing datasets. Altogether, 6768 GENCODE-annotated lncRNAs are represented across various compartments of 15 cell lines. We introduce “Relative concentration index” (RCI) as a useful measure of localisation derived from ensemble RNAseq measurements. LncATLAS is accessible through an intuitive and informative webserver, from which lncRNAs of interest are accessed using identifiers or names. Localisation is presented across cell types and organelles, and may be compared to the distribution of all other genes. Publication-quality figures and raw data tables are automatically generated with each query, and the entire dataset is also available to download.

**Conclusions:** LncATLAS makes lncRNA subcellular localisation data available to the widest possible number of researchers. It is available at lncATLAS.crg.eu.

## Introduction

The functions of long noncoding RNAs (lncRNAs) are intimately linked to location in the cell. The first discovered lncRNAs tended to be located in the nucleus and chromatin, and epigenetically regulate gene expression (Hutchinson et al. 2007; Mondal et al. 2010; Rinn et al. 2007; Tsai et al. 2010; Whitehead, Pandey, and Kanduri 2009; Zhao et al. 2008). However, we now appreciate lncRNAs’ localisation, and molecular functions, to be highly diverse. There exists a substantial population of lncRNAs in the cytoplasm (Carlevaro-Fita et al. 2016; van Heesch et al. 2014; Ulitsky and Bartel 2013), with evidence for roles such as translation regulation (Schein et al. 2016; Yoon et al. 2012; Zucchelli et al. 2016), miRNA decoys (Cesana et al. 2011), or protein trafficking (Aoki et al. 2010; Kino et al. 2010; Willingham et al. 2005). Consequently, ascertaining nuclear-cytoplasmic localisation has become one of the primary sources of evidence when investigating the molecular role of newly-discovered lncRNAs (J. Chen et al. 2016; L.-L. Chen 2016; Hansji et al. 2016; Hutchinson et al. 2007; Ishizuka et al. 2014; Ounzain et al. 2015).

The various methods to map RNA molecules in the cell operate with trade-offs in throughput, convenience, and accuracy. Amongst the single-gene approaches, probably the most commonly used is qRTPCR on RNA extracts of purified cellular compartments (Wang, Zhu, and Levy 2006). It yields information on relative RNA concentrations between compartments, but not of absolute molecule numbers per cell. Another method is fluorescence in situ hybridization (FISH), which can in principle yield absolute counts of molecules at subcellular resolution (Dunagin et al. 2015; Raj et al. 2008). However, FISH is time-consuming and low-throughput, and requires expensive reagents (Cabili et al. 2015). More recently, the ingenious in situ sequencing method, FISSEQ, has established high-throughput subcellular RNA counting (Lee et al. 2015). But at present just one dataset is available, and is restricted to several hundred highly-expressed lncRNAs (Lee et al. 2014).

The only method presently capable of whole-genome localisation mapping is subcellular RNA sequencing (subcRNAseq). Here cells are fractionated, and extracted RNA sequenced (Djebali et al. 2012). SubcRNAseq yields high-throughput and quantitative data, although as with RTPCR approaches mentioned above, the absolute counts of RNA molecules per cell are lost (Ulitsky and Bartel 2013). Recently, large amounts of raw subcRNAseq data have become available, most notably from the ENCODE consortium (Djebali et al. 2012; Dunham et al. 2012). These data remain under-utilized and have not been made readily accessible.

In light of the growing use of RNA localisation to infer function of newly-discovered lncRNAs, and the availability of large amounts of unprocessed subcRNAseq data, we have created a resource to make lncRNA localisation data available to the broader scientific community. This resource, “lncATLAS”, enables non-expert users to rapidly access a rich variety of easily-interpreted data on their lncRNA of interest.

## Results

### A database of lncRNA localisation based on human RNAseq data

In light of growing interest in lncRNA and their functions, we decided to create a resource for accessing and visualizing lncRNA localisation within human cells. We collected data from the largest dataset of subcRNAseq, produced by the ENCODE consortium (Djebali et al. 2012; Dunham et al. 2012). Raw RNAseq data from a panel of human cell lines were used to quantify the reference GENCODE gene annotation (Derrien et al. 2012; Harrow et al. 2012). RNAseq experiments were obtained for a total of 15 cell lines comprising 48 individual experiments (see Supplementary Table S1). These cells originate from a wide diversity of adult and embryological organ sites, and comprise both transformed and normal cells (Figure 1A). For each cell, cytoplasmic and nuclear data are available, and for the majority of these, whole-cell data were also obtained. In addition, from a single cell line, K562, sub-nuclear and sub-cytoplasmic compartment data are also available (Figure 1B; Supplementary Table S1). Hereafter we refer to these as “compartments”. PolyA+ RNA samples were available for whole cell, cytoplasm, nucleus and sub-cytoplasmic compartments, and total RNA for sub-nuclear compartments.

**Figure 1.**
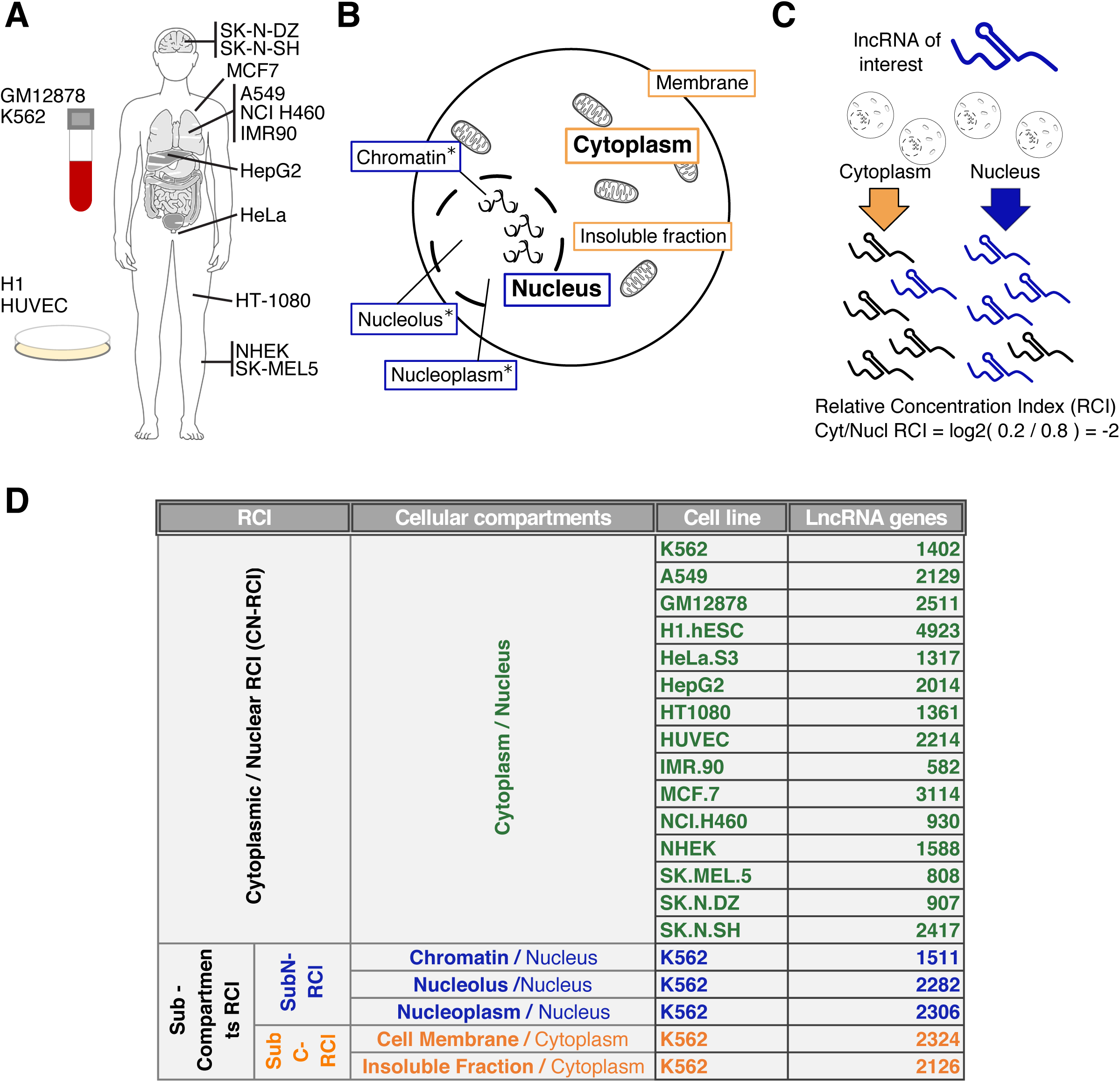
Overview of lncATLAS data. A) Cell lines available in lncATLAS, indicating their approximate origin. B) Cellular compartments available. * Compartments with only total RNA samples available. C) The relative concentration index (RCI), in this case calculated for the cytoplasm and nucleus (CN-RCI). The RCI can be thought of as the log-ratio, between two compartments, of the concentration of a given RNA molecule per unit mass of RNA. D) Overview of the cell lines and cellular compartments available for lncRNA RCI calculations. “Number of genes” indicates the number of lncRNA for which the RCI could be calculated in the corresponding cell line (see Materials and Methods for details). Sub-C RCI and Sub-N RCI correspond to sub-cytoplasmic RCI and sub-nuclear RCI respectively.

### Defining localisation from RNAseq data

Throughout the present study, for practical reasons, we adopt a relative scheme to define and quantify RNA localisation: the “Relative Concentration Index” (RCI). RCI is defined as the log2-transformed ratio of FPKM (fragments per kilobase per million mapped) in two samples, for example the cytoplasm and nucleus (Figure 1C). A similar approach has been used previously (Derrien et al. 2012; Ulitsky and Bartel 2013). It is worth commenting on exactly how these values should be interpreted: RCI is the ratio of a transcript’s concentration, per unit mass of sampled RNA, between two compartments. Sampled RNA populations may be PolyA+ RNA or total RNA, and we are careful to only compute RCI values within the same population.

The mass of RNA per compartment per cell is not equal, and typically not quantified prior to RNAseq (Djebali et al. 2012). Therefore, without knowing the total mass of PolyA+ RNA in the nucleus and cytoplasm of a single cell, we cannot make statements about the relative *number* of RNA transcripts in cellular compartments of a single cell (Cabili et al. 2015).

We here briefly digress to contrast this approach with another possible way to define relative subcellular localisation. Perhaps more obvious is a “molecular” definition, in terms of numbers of molecules of a given RNA transcript *X* in the compartments of a single cell. For example, if one cell has 10 and 5 molecules of *X* in the cytoplasm and nucleus, respectively, then its cytoplasmic/nuclear localisation would be defined as 10/5=2. We define this measure as “Relative Molecules Index” (RMI). Such information is, in principle, directly accessible from fluorescence-based techniques, and has been calculated previously (Cabili et al. 2015; Lee et al. 2014). As mentioned above, our ignorance of the total PolyA+ RNA mass of the cell lines used here, precludes the calculation of RMI in this study.

### Computing localisation across genes and cell types

RCI was calculated for various selected pairs of cellular compartments (Figure 1D). In the majority of cases, we calculated the cytosoplasmic/nuclear RCI – “CN-RCI” (Supplementary Table S2) (see Materials and Methods). This is a measure of the relative concentration of an RNA sequence in the cytoplasm, compared to the nucleus, in log2 units. For one cell type, K562, total RNA data from sub-nuclear and PolyA+ RNA from sub-cytoplasmic compartments were also calculated, by reference to total RNA from the nucleus or PolyA+ RNA from the cytoplasm, as appropriate (Supplementary Table S3) (see Materials and Methods). Altogether this yielded localisation estimates in 20 compartment / cell combinations (Figure 1D).

Where available, replicate data were used to assess reliability of RCI measurements (see Supplementary Table S1 for information about availability of replicates). Silent and unreliable genes were excluded from further consideration (see Materials and Methods for details in the filtering steps). Between 3114 (max) and 582 (min) lncRNAs’ CN-RCI localisation could be estimated per cell, after filtering (Figure 1D,2A). Note that H1.hESC has a greater number of detected genes (4923 genes), because biological replicates were not available for cytoplasm nor nucleus for this cell line. A total of 24,538 genes (17,770 mRNAs and 6,768 lncRNAs) were quantified in at least one cell type. Of these, 31 lncRNAs were detected in all samples tested (Figure 2B and Supplementary Table S4). LncRNAs display a highly cell-type specific detection pattern, in contrast to mRNAs, as observed previously (Figure 2B) (Cabili et al. 2015; Derrien et al. 2012; Guttman and Rinn 2012).

RCI data are consistent with known cytoplasmic-nuclear localisation tendencies of lncRNAs and mRNAs. Amongst the top 15 most cytoplasmic measurements, 14 represent mRNAs (the remainder is an annotation of uncertain biotype and may be protein-coding) (Supplementary Table S5). In contrast, 12 of the 15 most nuclear RCI values represent lncRNAs (Supplementary Table S6). The nuclear-enriched X-chromosome inactivating transcript *XIST* occupies the top four positions (Brown et al. 1992; Clemson et al. 1996). Manual inspection of several well-known lncRNAs showed that localisation reported here tended to be consistent with literature reports (see next Section).

Detected lncRNA genes reported by lncATLAS cover a substantial fraction of the entries from manually-curated and widely-used databases, such as lncRNAdb and LncRNADisease, and new localization specific databases such as RNALocate (Zhang et al. 2016). Note that these databases contain a mixture of GENCODE annotated and non-GENCODE annotated entries. 74, 128 and 150 genes from lncRNAdb, lncRNADisease and RNALocate respectively are detected in lncATLAS. These numbers represent a 39%, 48% and 47% of the total number of human lncRNAs and a 63%, 73% and 90% of the total number of human GENCODE annotated lncRNAs of each database, respectively (Figure 2C). LncRNAs from these databases that are not displayed in lncATLAS are either not detected in any of the cell lines considered, or else do not belong to the GENCODE annotation.

**Figure 2.**
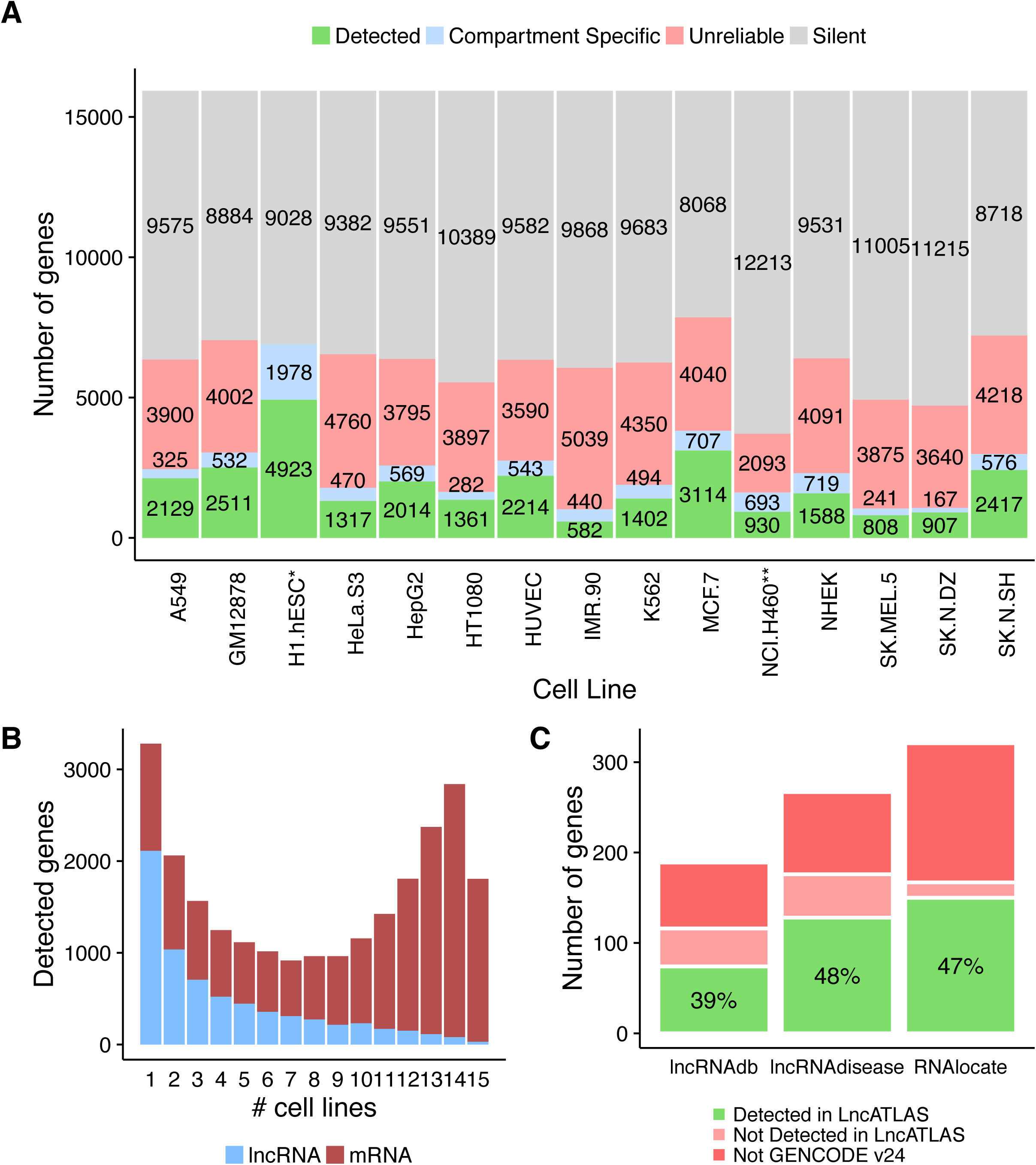
Summary of Cytoplasmic/Nuclear detected genes by lncATLAS. A) Number of genes analysed in lncATLAS. (i) Detected genes: present a reliable expression value in both compartments, cytoplasm and nucleus. Other categories comprise (ii) Compartment specific genes: present a reliable non-zero value in one compartment and zero in the other, (iii) unreliable genes: genes that did not pass the 2fold cutoff (see Materials and Methods, data processing) and (iv) silent genes: not expressed in any compartment (see Materials and Methods, CN-RCI). *No biological replicates were available for cytoplasm nor nucleus. **No biological replicates were available for cytoplasm. B) Histogram showing how many genes are detected in a determined number of cell lines. LncRNA genes in blue and mRNAs in red. C) Coverage by lncATLAS of widely-used, manually-curated lncRNA databases, lncRNAdb (Quek et al. 2015) and lncRNAdisease (Chen et al. 2013). The new RNAlocate database (Zhang et al. 2016) is also shown for comparison. Note that these databases hold a mixture of GENCODE annotated and non-GENCODE annotated lncRNAs. The barplot displays the total number of human lncRNA genes in each database (whole bar). Bars are colored to represent, from the total number of human lncRNAs in a database, the number of genes that: (green) are displayed in lncATLAs (the percentage numbers indicate the proportion that this fraction represents), (shaded red) are part of GENCODE annotation but are not detected in any of the 15 cell lines, and therefore are not displayed in lncATLAs, (red) are not present in GENCODE annotation and could not be considered in our database.

### LncATLAS webserver for exploring localisation data

The lncATLAS dataset was compiled into a relational database that is searchable through a webserver at lncATLAS.crg.eu. LncRNAs of interest are accessed using official gene names or GENCODE gene identifiers. A maximum of three genes may be investigated simultaneously. Several well-known lncRNAs with known localisation are also available for reference. Once a gene or genes has been selected, a series of data interpretations are presented, summarized below. As examples, results for *MALAT1* (nuclear localized) (Hutchinson et al. 2007) and *DANCR* (cytoplasm localized) (Cabili et al. 2015; van Heesch et al. 2014) lncRNAs data are shown in Figure 3 and 4.

The following sections summarise the data presented to a user for their gene of interest (GOI).

*1. Inspect the cytoplasmic-nuclear localisation of your gene of interest (GOI*)

Data are only displayed for selected gene(s).

**Plot 1: Cytoplasmic/Nuclear Localisation: RCI and expression values (all cell types):** In this basic summary, the Cytoplasmic/Nuclear RCI is shown as a bar plot across all available cell types. Bars are coloured to reflect the expression level of the gene, as inferred from nuclear RNAseq. The individual FPKM values, upon which RCI values are based, are displayed. When a gene is expressed only in one compartment, RCI cannot be computed; then, dashed bars with expression values are shown instead (Figure 3A).

*2. Inspect the cytoplasmic-nuclear localisation of your GOI within the distribution of all genes*.

The aim of the second section is to understand, in terms of localisation, how the genes of interest behave relative to all other genes. Three different plots show CN-RCI values distribution for all lncRNAs and mRNAs, within which the location of the GOI is indicated.

**Plot 2: Cytoplasmic/Nuclear Localisation: RCI distribution (all cell types):** To put RCI values in context, their percentile rank within the distribution of all lncRNAs is indicated (ranks relative to lowest value). Data are shown for all cell types (Figure 3B).

**Plot 3: Cytoplasmic/Nuclear Localisation: RCI distribution (individual cell type):** The same data are shown as for Plot 2, but in the form of a density plot. The User must here specify a single cell type. When genes are not classed as “Detected”, RCI cannot be computed and no data are shown (Figure 3C).

**Plot 4: Cytoplasmic/Nuclear Localisation: Comparison with expression (individual cell type):** As for Plot 3, gene values are shown in the context of all other genes in the same cell, but here also indicating whole-cell expression values. As before, the data are shown for a single cell type chosen by the User, and plots are only generated for cells where RCI values are “Detected” (Figure 3D).

*3. Inspect the localisation of your GOI at sub-compartment level*.

The final section gives information about enrichment in the cytoplasmic and nuclear subcompartments of K562 cells. As in the previous section, RCI values for the genes of interest are indicated in the context of full lncRNA and mRNA distributions.

**Plot 5: Sub-cytoplasmic and Sub-nuclear Localisation in K562 Cells:** Here data are shown for sub-nuclear and sub-cytoplasmic compartments K562 cell line. As in Plots 2 and 3, distributions across all detected genes are shown (Figure 4).

In the examples shown, the differences in localisation of *MALAT1* and *DANCR* are clear. Their cytoplasmic-nuclear localisations are highly divergent (Figure 3 and 4), and broadly consistent across all the cell lines observed. The difference in localisation is observed even in cells where their overall expression level is similar (eg HUVEC, Figure 3D).

All figures may be downloaded as publication-quality files in pdf format. Similarly, in the *Get Raw Data* tab, the underlying RCI and raw expression values for selected genes may be accessed as a batch query. Furthermore, the entire set of data tables for lncATLAS may be downloaded in the same tab using the *Download All raw data* button.

**Figure 3.**
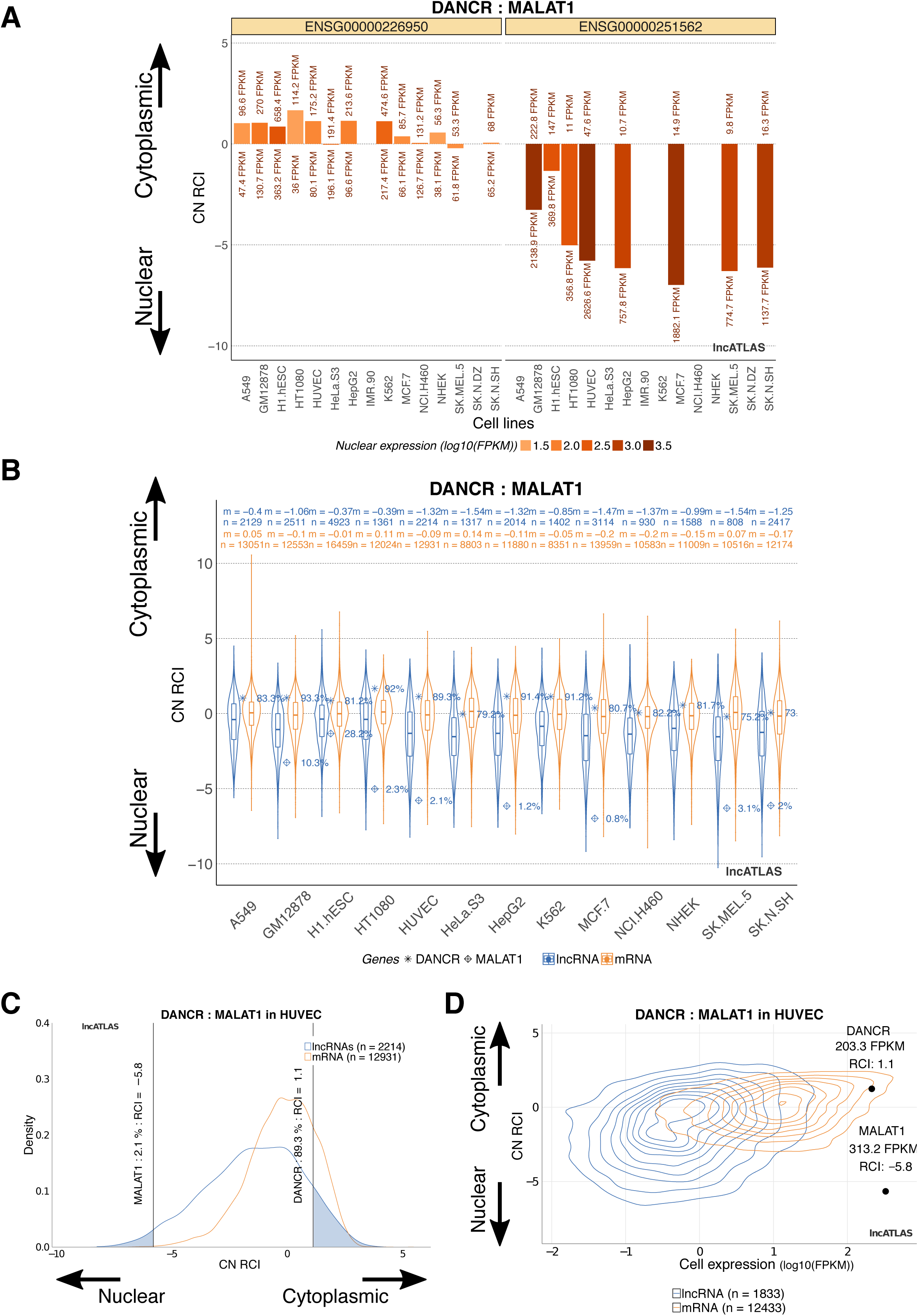
Subcellular localisation plots displayed by lncATLAS. MALAT1 and DANCR genes are selected as examples of nuclear and cytoplasmic lncRNAs, respectively. A) Bars representing CN-RCI values for the selected genes across all cell lines. Expression values (FPKMs) for the genes of interest are shown for both compartments (cytoplasm on top of the bar, nucleus on the bottom). Bars are colored by their absolute nuclear expression. B) Boxplot showing CN-RCI values distribution of all lncRNAs (blue) and mRNAs (orange) for each cell line (“n” indicates total number of genes, “m” median of CN-RCI values for lncRNAs and mRNAs separately). LncRNAs of interest are located in the distribution and a percentage indicates their percentile rank within the distribution of all lncRNAs (ranks relative to lowest value). C) Same than in the previous plot but in this case distribution is shown as a density plot and only for a particular cell type, HUVEC. Again, genes of interest are located in the distribution and their percentile rank (relative to lowest value) and RCI are indicated. D) Contour plot showing lncRNA and mRNA populations as a function of CN-RCI values and whole cell expression (log10(FPKMs)). LncRNAs of interest are specifically displayed together with their whole cell expression and RCI.

**Figure 4.**
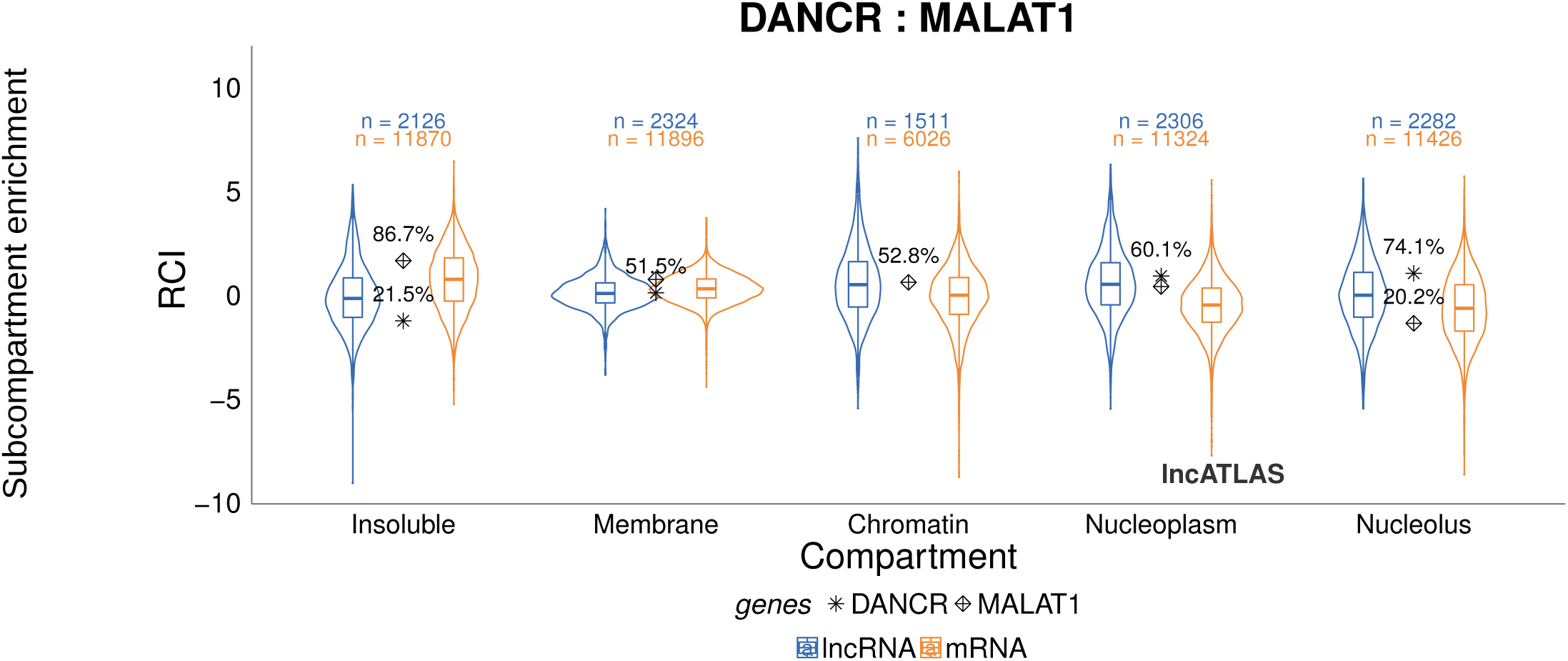
Sub-compartment data as displayed in LncATLAS. For K562 cells subnuclear and subcytoplasmic fractions were available and RCI for all lncRNAs and mRNAs was computed (see Materials and Methods). Boxplot shows the distribution of these values for each subnuclear and subcytoplasmic compartment (“n” indicates total number of genes in a distribution). Percentile rank (relative to lowest value) of each gene of interest is displayed to contextualize the relative enrichment of these genes in a subcompartment compared to the rest of lncRNAs and mRNAs.

## Discussion

Subcellular localisation provides important clues to the molecular function of novel lncRNAs. LncATLAS is designed to make such data available to the largest number of researchers. To our knowledge, only one other database of lncRNA localisation exists: RNALocate (Zhang et al. 2016). RNALocate contains manually-curated localisation classifications across multiple species. Despite focussing on a single species (human), due to the limited availability of subcellular RNAseq data, LncATLAS has two key advantages: it is quantitative, and it is based on standard GENCODE annotations, the *de facto* official annotation for both protein-coding and lncRNA genes (Derrien et al. 2012). These features boost the usefulness of lncATLAS data for other research groups and ensure its integration with diverse other genomics datasets. Future subcellular RNAseq data from other cell types, or other species, will be integrated as they become available.

## Materials and Methods

### Data source

Cytoplasmic and nuclear PolyA+ RNAseq data from 15 different cell lines were obtained from ENCODE (Djebali et al. 2012). (ENCODE RNAseq data in BAM format were obtained from ENCODE Data Coordination Centre (DCC) in September 2016 - https://www.encodeproiect.org/matrix/?type=Experiment). For most cell lines, whole-cell data were also obtained (exceptions being HT1080, NCI.H460, SK.MEL.5 and SK.N.DZ). A full list of processed RNAseq libraries are available in Supplementary Table S1.

### Data processing

Data were mapped to human genome assembly GRCh38 using STAR software (Dobin et al. 2013) and quantified with RSEM (Li and Dewey 2011) for all GENCODE v24 gene quantification, within the GRAPE analysis pipeline (Harrow et al. 2012; Knowles et al. 2013). Data consisted of two independent biological replicates per cell line and fraction (exceptions being H1.hESC cytoplasm and nucleus and NCI.H460 cytoplasm for which only one replicate was available) (see Supplementary Table S1 for a full list of source datasets). For sub-cytoplasmic RCI, instead of using poly+ cytoplasmic sample coming from Gingeras lab (used for CNRCI), we used the corresponding sample from the lab where the sub-cytoplasmic fractionation was done (Lécuyer lab). This is not considered as an additional biological replicate.

Throughout, RNAseq data are processed at the level of genes, rather than transcripts. From the whole GENCODE v24 annotation, genes contained in the “Long non-coding RNA gene annotation” define our lncRNA set of genes. Protein coding gene set is defined by GENCODE biotype “protein_coding” (see Supplementary Table S7).

In order to remove genes with high variability between replicates, genes with >2-fold difference between replicates are labelled “unreliable” and excluded from further analysis. This cutoff was not possible for the samples, mentioned above, for which replicate experiments were not available.

### Cytoplasmic-nuclear relative concentration index (CN-RCI)

For cytoplasmic/nuclear localisation, PolyA+ RNA data were used. At this stage, all genes are defined in one of four categories in each cell line: (1) Detected: genes with non-zero and “reliable” values in both cellular compartments; (2) Compartment-specific: genes considered “reliable” in both compartments but expressed >0 FPKM only in one; (3) Silent: both compartments have FPKM=0; (4) Unreliable: genes that are unreliable in at least one compartment.

Genes classed as “Detected” were retained, and their localisation was computed as the C/N Relative Concentration Index (CN-RCI) thus:

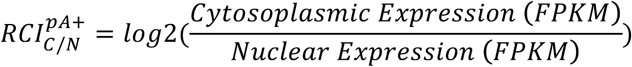

Only detected genes are shown in plots of lncATLAS, with the exception of compartment-specific genes in Plot 1. For this group of genes, CN-RCI value is not available in the plot and the bar only indicates the tendency of the gene towards nucleus or cytoplasm. Colour and FPKM are shown normally, to indicate the level of expression.

### Sub-cytoplasmic and sub-nuclear fractions for K562 (subN RCI, subC RCI)

RNAseq data from subnuclear fractions (chromatin, nucleolus and nucleoplasm) and from sub-cytoplasmic fractions (membrane and insoluble fraction) were retrieved and processed as above. These data are only available for K562 cells, and in the case of subnuclear samples correspond to total RNA (not PolyA+ selected RNA). Data were processed and RCI calculated as above, with the only differences being: (1) the RCI was calculated with reference to the nuclear fraction for sub-nuclear compartments, and cytoplasmic fraction for sub-cytoplasmic compartments; (2) for subnuclear compartments total RNA samples were used, instead of PolyA+.

### Database design

LncATLAS is a relational database implemented in MySQL (http://www.mysql.com) and designed through the official MySQL WorkBench tool for Linux. The Entity-Relationship (ER) diagram extracted summarizes its structure (Supplementary Figure S1). The tables are hierarchically organized from general information of the samples to the expression value per gene that is stored in the expression table.

### Web-tool Implementation

LncATLAS is constructed using the Shiny R package (version 0.13.2) (http://www.rstudio.com/shiny/). The database is connected to the application itself via the R package RMySQL (version 0.10.9). Other packages used in the implementation are the *ggplot2* package (version 2.1.0) and *dplyr* (version 0.5.0) used to build custom plots and manipulate the data.

## Acknowledgements

We acknowledge ENCODE Consortium and the respective laboratories producing the data. We also acknowledge support of the Spanish Ministry of Economy and Competitiveness, ‘Centro de Excelencia Severo Ochoa 2013-2017’, SEV-2012-0208. R.J. was supported by Ramón y Cajal RYC-2011-08851. This research was partly supported by the NCCR RNA & Disease funded by the Swiss National Science Foundation.

